# Investigation of Chordwise Functionally Graded Flexural Rigidity in Flapping Wings Using a Two-Dimensional Pitch-Plunge Model

**DOI:** 10.1101/2021.08.27.457983

**Authors:** Joseph Reade, Mark Jankauski

**Affiliations:** Montana State University, Dept. of Mechanical & Industrial Engineering, Bozeman, MT

**Keywords:** flapping wings, unsteady vortex lattice method, flight energetics, reduced-order modeling, pitching-plunging airfoil

## Abstract

Insect wings are heterogeneous structures, with flexural rigidity varying one to two orders of magnitude over the wing surface. This heterogeneity influences the deformation the flapping wing experiences during flight. However, it is not well understood how this flexural rigidity gradient affects wing performance. Here, we develop a simplified 2D model of a flapping wing as a pitching, plunging airfoil using the assumed mode method and unsteady vortex lattice method to model the structural and fluid dynamics, respectively. We conduct parameter studies to explore how variable flexural rigidity affects mean lift production, power consumption and the forces required to flap the wing. We find that there is an optimal flexural rigidity distribution that maximizes lift production; this distribution generally corresponds to a 3:1 ratio between the wing’s flapping and natural frequencies, though the ratio is sensitive to flapping kinematics. For hovering flight, the optimized flexible wing produces 20% more lift and requires 15% less power compared to a rigid wing but needs 20% higher forces to flap. Even when flapping kinematics deviate from those observed during hover, the flexible wing outperforms the rigid wing in terms of aerodynamic force generation and power across a wide range of flexural rigidity gradients. Peak force requirements and power consumption are inversely proportional with respect to flexural rigidity gradient, which may present a trade-off between insect muscle size and energy storage requirements. The model developed in this work can be used to efficiently investigate other spatially variant morphological or material wing features moving forward.

## 1. Introduction

Flapping insect wings are highly compliant structures that deform under aerodynamic and inertial forces [1]. Deformation is believed to play a fundamental role in insect flight mechanics and has been shown to enhance aerodynamic force generation [2–4] and energetic efficiency [5–7]. Various structural features, including venation and corrugation, influence wing deformation under dynamic loading. These features also govern how the wing’s mechanical properties vary in space. Experimental studies have shown that the flexural rigidity of the Hawkmoth *Manduca sexta* forewing, for example, varies one to two orders of magnitude over the wing’s chord length [8]. This pronounced gradient may underpin wing responses that are aerodynamically and energetically favorable. However, it is presently not well understood how graded flexural rigidity influences flapping wing performance.

This is in part because models used to predict wing fluid-structure interaction (FSI) are computationally-intensive. They are consequently challenged by parameter studies that consider variable flexural rigidity or other morphological and material properties. Flapping wing FSI models usually rely on coupled finite element method (FEM) and computational fluid dynamic solvers (CFD) [9–13]. Flapping wings pose unique challenges to both of these numerical approaches. Given the finite, periodic rotation of the wing, stiffness matrices within FEM must be updated at each time interval of analysis, which increases computation times considerably [14]. CFD requires that the Navier-Stokes equations be solved across a discretized fluid domain [15], which could result in upwards of hundreds of thousands of equations to solve. Estimating wing deformation via coupled CFD/FEM may therefore span several several hours per wingbeat in some cases.

Lower-order FSI models can decrease the computational requirements considerably. Simplifications in fluid and structural solvers, as well as converting the 3D problem to a 2D idealization, have yielded models that can be solved on the order of seconds. Some of the most common flapping wing FSI models treat the wing as a 2D pitching, plunging airfoil [16, 7, 17, 2, 3]. The airfoil represents a cross-section running along the chord of the insect’s wing. Plunge represents the primary flap rotation and pitch represents the rotation the wing experiences about its leading edge. The 2D pitch-plunge framework neglects some of the physics that play a role in true 3D flapping, such as spanwise bending [18], spanwise flow [19] and wingtip effects [20]. Despite lower quantitative accuracy relative to 3D models, 2D models can quickly identify solution trends and thus can establish foundational knowledge that may be subsequently extended to more computationally intensive 3D modeling. For this purpose, they have been widely utilized to investigate several aspects of flapping wing mechanics. For example, Yin and Luo used a pitch-plunge model to show that, when a wing’s deformation was dominated by external flows rather than inertia, it had better power efficiency [7]. Tian et al. used this framework to understand how asymmetry between the wing’s deformation during upstroke and downstroke affected forward flight [17]. Vanella et al. demonstrated that a super-harmonic resonance may favorably influence the wing’s lift to drag ratio at low Reynold’s numbers [2]. Mountcastle and Daniel investigated how variable wing kinematics and structural properties affected the wing’s lift generation [3].

Despite the contributions of these works and others, the influence graded flexural rigidity has on flapping wing aeromechanics remains understudied. The objectives of the present research are to (1) develop a reduced-order FSI model of a flapping, flexible insect wing treated as a pitching plunging airfoil, and (2) to use this model to understand how graded flexural rigidity affects wing dynamics. Specifically, we evaluate lift, power and the forces required to drive the wing. We employ a number of assumptions to simplify the wing structure and flapping kinematics in this study. First, because we treat the wing as a 2D pitching, plunging airfoil, spanwise flow, spanwise bending and wingtip losses associated with 3D motion are neglected. Second, we neglect complex venation patterns, cross-sectional curvature and spatially-variable material properties that may affect the wing’s local flexural rigidity. We instead assume that spatial variation of flexural rigidity is driven by local variations of effective wing thickness, since vein thickness reduces dramatically along the chord in many insect wings [21]. Because of these assumptions, the model developed in this work is not suitable for high-fidelity fluid or structural calculations of real insect wings. It is instead intended to isolate and study variable flexural rigidity in a simpler setting.

The remainder of this manuscript is organized as follows. First, we derive the FSI framework using the assumed mode method (AMM) to model the structure and unsteady vortex lattice method (UVLM) to model the fluid. Next, we apply the model to numerically simulate the response of a wing with similar properties to a Hawkmoth *Manduxa sexta* (*M. sexta*) forewing. We explore the aerodynamics and energetic requirements of the wing under hovering flapping circumstances, and conduct a parameter study to determine how these quantities are influenced by variable kinematics. We conclude by discussing the broader implications of this research on the study of insect flight.

## 2. Theory

### 2.1. Structural Model

In this section, we derive the equation of motion governing the wing’s deformation via the Lagrangian approach. This structural model was derived in [22] and is summarized here for clarity. We treat the wing as an Euler-Bernoulli beam, where the leading edge is clamped and the trailing edge is free to deflect. Wing deformation is calculated using the AMM, where the total deflection is a linear combination of the wing’s eigenfunctions.

First, we first define two reference frames; an inertial *X* – *Y* – *Z* frame that is fixed in space, and a body-fixed *x* – *y* – *z* frame that translates and rotates with the wing’s rigid body motion (Fig. 1). Note that the gravity would act in the +Y direction, and thus the wing must produce lift in the -Y direction in order for the insect to remain aloft. The origin *O* of the *x* – *y* – *z* frame is located at the leading edge of the wing. *O* is subject to prescribed plunging *Z*(*t*) and the wing experiences prescribed pitching *θ*(*t*) about *O*. The *x* – *y* – *z* frame has angular velocity

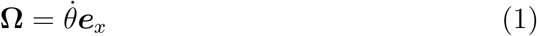

**Figure 1:**
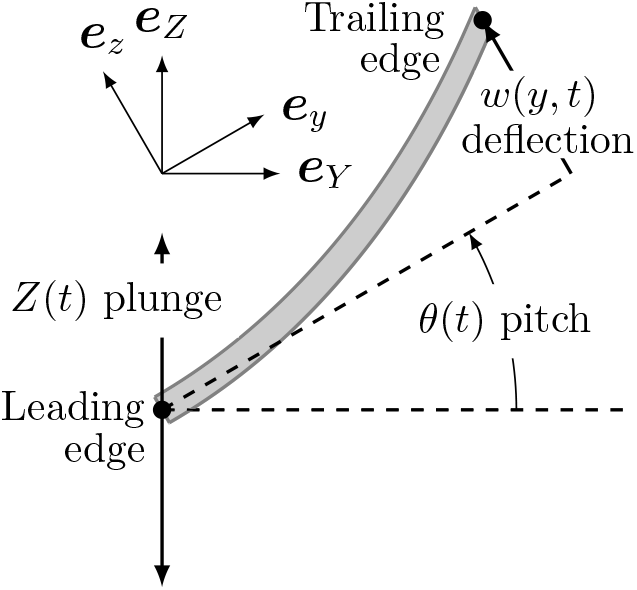
Reference frames and pitch-plunge motion.

The position ***R*** of a differential mass element *dm* relative to the wing-fixed frame is the sum of three component vectors (e.g., ***R*** = ***r***_1_ + ***r***_2_ + ***r***_3_), where

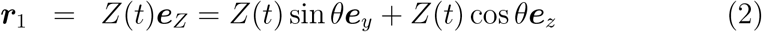

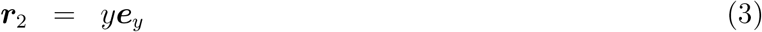

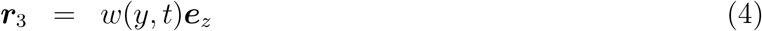

Above, ***r***_1_ describes the displacement associated with the plunging motion, ***r***_2_ describes the position of dm at a location *y* along the wing’s chord, and ***r***_3_ describes an infinitesimal out-of-plane wing deflection *w*. In-plane deflection is neglected. The velocity of *dm* is

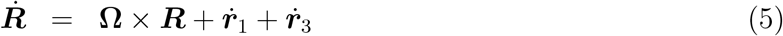

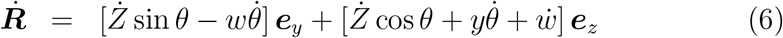

and the acceleration of *dm* is

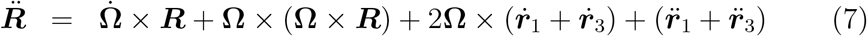

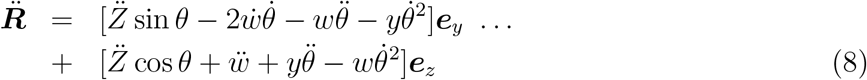

While 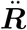 is not required to derive the equation of motion, it is necessary to calculate power consumption and inertial forces and moments as detailed later. We expand wing deflection *w*(*y,t*) in terms of mass-normalized mode shapes *ϕ_k_* and modal responses *q_k_* as

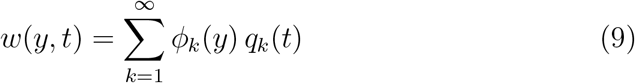

The wing’s total kinetic energy *T* and potential energy *U*

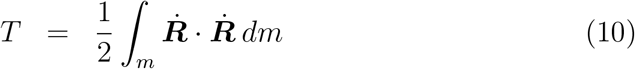

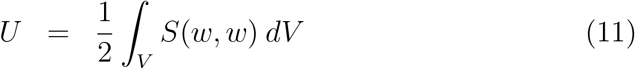

where *S* is a symmetric, quadratic strain energy density function and *V* is the wing’s volumetric domain. Applying Lagrange’s equation, we arrive at the equation of motion governing modal response *q_k_* as

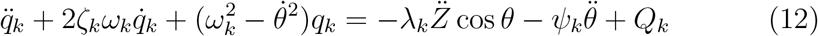

where *ω_k_* and *ζ_k_* are the wing’s k^*th*^ natural frequency and damping ratio, respectively. Note that the viscous damping term is not derived explicitly through the energy formulation and is instead included as an empirical correction term. Constants λ_*k*_ and *ψ*_*k*_ are obtained by integrating the mode shape over the wing’s mass domain, or

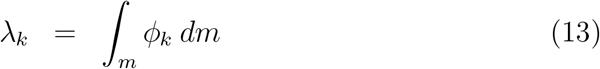

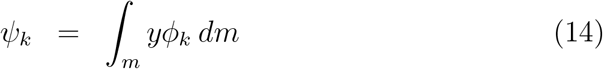

Eq. 12 shows that the wing’s stiffness is time-varying and dependent on the wing’s pitching rate and stationary natural frequency. The first excitation term to the right hand side of Eq. 12 is a plunge-dominated inertial term proportional to the linear acceleration of the wing. The second excitation term is a pitch-dominated inertial term modulated by the wing’s angular acceleration. The non-conservative generalized load *Q_k_* is obtained by integrating the projection of an aerodynamic force distribution onto the mode shape

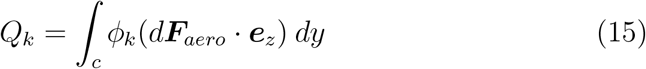

where *d****F***_*aero*_ is the aerodynamic force per unit length, discussed the following section.

### 2.2. Fluid Model

We use a modified UVLM approach to predict the flow and estimate the force distribution over the wing. Our method is based on that described in [23]. The UVLM is based on potential flow theory. The flow is assumed to be irrotational, except on the surface of the wing and in the wake that is shed from the trailing edge, and is incompressible throughout. By modeling the wake, we account for unsteady effects such as added mass. Viscous effects are not accounted for.

The wing is discretized into *N_p_* panels, each of length *ds* (Fig. 2). On each panel, a bound vortex and control point are located 0.25*ds* and 0.75*ds* from the leading edge of the panel, respectively [24]. Additionally, a wake vortex is shed from the trailing edge of the wing at each time step, with the wake being truncated at *N_w_* vortices.

**Figure 2:**
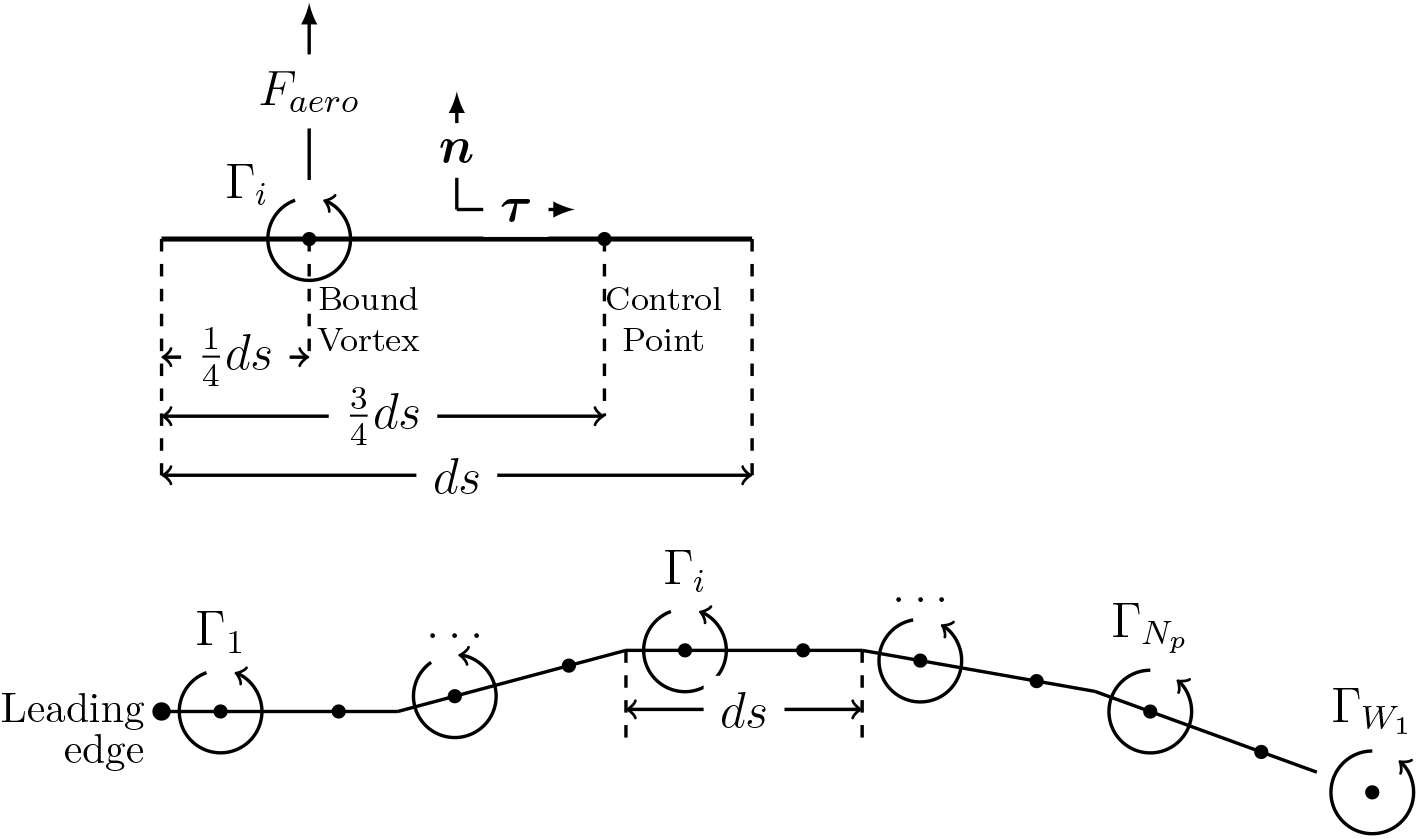
Wing discretization used by UVLM.

**Figure 3:**
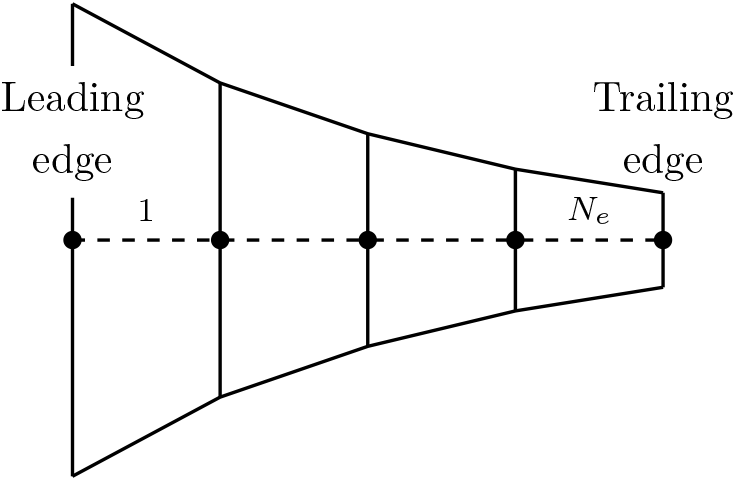
Wing discretization with exponential thickness distribution. Nodes 2,…, *N_e_* + 1 may experience vertical displacement and rotation, while the leading-edge node is fixed. Wing thickness is exaggerated.

The bound vortex induces a flow velocity around itself that diminishes with distance. The flow velocity at any point *i* in space that is induced by a vortex at *j* is given by the Biot-Savart law:

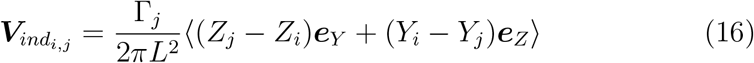

where ***V***_*ind*_ is the induced velocity, *L* is the distance between point *i* and vortex *j, Y* and *Z* are the coordinates of *i* and *j* in the inertial reference frame, and Γ_*j*_ is the circulation of the vortex. The Kelvin-Helmholtz theorem for an ideal fluid states that the total circulation around a closed contour must remain constant, or

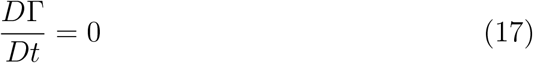

Since the initial state of the fluid flow is at rest with zero vorticity throughout, this implies that the total vorticity, found by summing the strengths of the bound and wake vortices, is zero at each time step. This condition is used to calculate the strength of the wake vortices.

The non-penetration boundary condition states that the flow may not travel through the solid surface of the wing, meaning that the component of velocity normal to the surface is zero. This is represented by

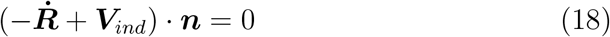

where 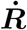 is the velocity of the wing’s surface in the inertial reference frame and ***n*** is the surface normal vector. Because there are a finite number of bound vortices, the non-penetration condition cannot be satisfied across an entire panel, so the control point is chosen as the location where it must hold true. By simultaneously enforcing the non-penetration condition at all control points on the wing, as well as the Kelvin-Helmholtz condition, the strengths of the bound vortices can be determined via

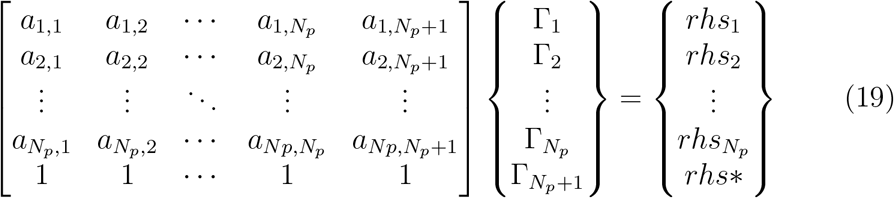

where *a_i,j_* is the influence coefficient. On the right-hand side, *rhs*_1,2_…*N_p_*, are the normal velocities due to the combined effect of the pitch-plunge motion, structural deflection, and velocity induced by the wake vortices. *rhs*\* is the total bound vorticity from the previous time step. The solved quantities Γ1,2… *N_p_* and Γ*N_p_*+1 are the strengths of the bound vortices on the wing and of the newly-shed wake vortex, respectively. The new wake vortex is placed a fixed distance aft of the trailing edge. All existing wake vortices are advected with the local flow. The velocity of the *i^th^* wake vortex is given by

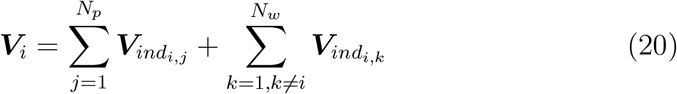

where ***V***_*i*_ is the total induced velocity at *i* by *N_p_* bound vortices on the wing and *N_w_* – 1 wake vortices. A wake vortex is not influenced by its own vorticity. The aerodynamic force on the wing is determined using the unsteady Bernoulli equation,

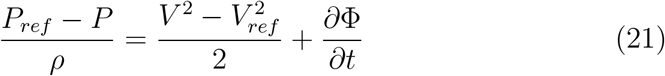

where *P* and *P_ref_* are the local and reference pressures, *V* and *V_ref_* are the local and reference (freestream) velocities, *ρ* is the fluid density, and Φ is the flow potential. When solved at the bound vortex for the *i^th^* wing panel, this becomes

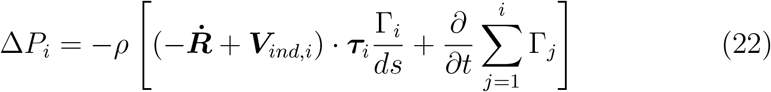

where Δ*P* is the pressure difference between the top and bottom surfaces of the wing, ***V***_*ind*_ is the velocity induced by the bound and wake vortices, and *τ* is the unit vector tangent to the wing surface. The total aerodynamic force is obtained by summing the products of the pressure differentials and the panel normal vectors.

### 2.3. Force, Moments and Power

Several quantities of interest are not directly calculated by the AMM or UVLM, such as the power required to flap the wing or the forces and moments necessary to maintain the prescribed motion of *O*. Flight is the most energetically expensive mode of locomotion per unit of time, and thus it is essential to understand how the wing’s structural dynamics underpin low energetic expenditures. In this section, we derive expressions based on modal responses to calculate flapping wing moments, forces and instantaneous power.

Total power consumption is broken into two components: inertial power and aerodynamic power. Inertial power is required to move the wing mass through space - if the wing was operated in a vacuum, inertial power would equal to total power. Inertial power of a rotating, translating system is

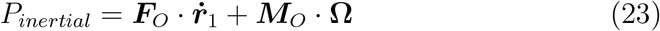

where ***F***_*O*_ is the inertial force at *O* and ***M***_*O*_ is the inertial moment about the leading edge. The inertial force is the wing’s spatiotemporal acceleration integrated over the wing’s mass, given by

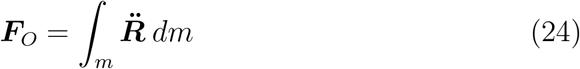

When Eq. 8 is substituted in for 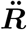, this becomes

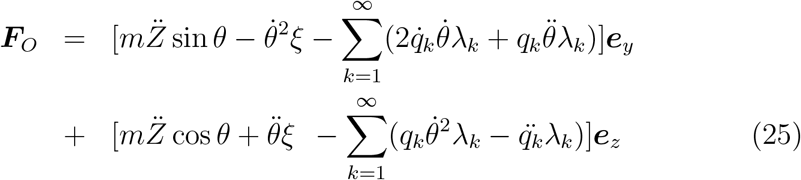

where *m* is the mass of the wing and *ξ* is a constant defined by

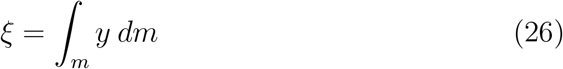

The inertial moment about *O* is

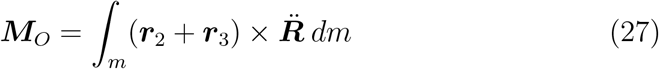

When treated in the same manner as the inertial force, this expression becomes

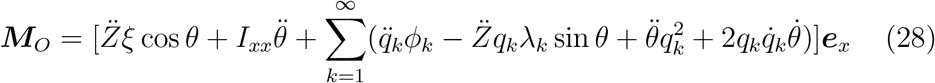

where *I_xx_* is the wing’s mass-moment of inertia about *O*. The aerodynamic power is the power required to overcome the fluid forces acting against the wing, given by

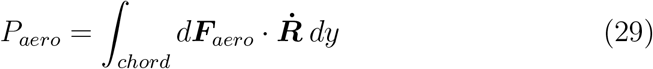

where *d****F***_*aero*_ is the aerodynamic force at some point on the wing as computed by the UVLM.

## 3. Numerical Simulation

Here, we describe the numerical parameters used to simulate the dynamic response of the wing. We use a finite element approach to convert the wing from a continuous domain to a discrete domain. The wing is discretized into *N_e_* beam elements each with two degrees-of-freedom (vertical displacement and rotation) at each node. The wing thickness decays exponentially from leading edge to trailing edge, with the thickness *t* at chordwise location *y* described by

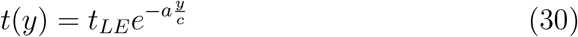

where *t_LE_* is the leading edge thickness, *c* is the chord length and *a* is the decay rate. Values of *a* range from 0 to 1.6, which correspond to leading-to-trailing-edge taper ratios *t_TE_/t_LE_* from 1.0 (uniform thickness) to 0.2 (highly tapered). Rigid wings are simulated for comparison. The most tapered case results in a two orders-of-magnitude decrease in flexural stiffness over the chord, which encompasses the range of chordwise-stiffness distributions reported in *Manduca sexta* [8].

The stiffness matrix **K**_*e*_ based on Euler-Bernoulli beam theory for an individual element is

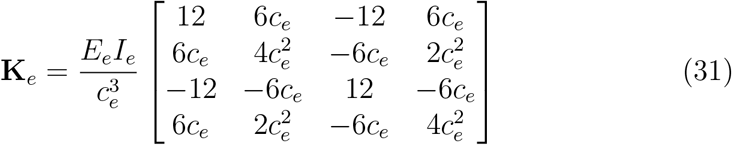

where *I_e_, E_e_*, and *c_e_* are the elemental area moment of inertia, elastic modulus, and length respectively. In the *I_e_* calculation, the cross-sectional area of the wing in the *x* – *z* plane is assumed rectangular. *E_e_* and *c_e_* are uniform across all elements. The elemental mass matrix **M**_*e*_ is

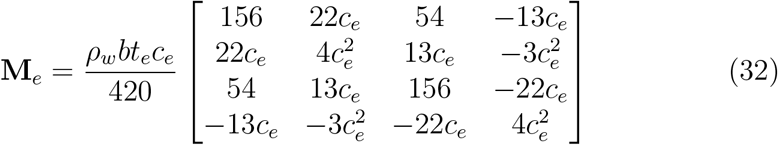

where *ρ_w_* is the wing material density, *b* is the wing span, and *t_e_* is the element thickness.

Discretization parameters are summarized in Tab. 1. Peak wing deflection converged at 20 panels in the UVLM solver. The most costly quantity in the UVLM was the number of wake vortices, as this was far greater than the number of wing panels and the calculations per step is proportional to (*N_p_* + *N_w_*)^2^. 40 panels were used in all simulations to account for variation in the conditions. While the fundamental frequency of the beam converged within 20 elements for the most tapered case, for simplicity *N_e_* is set equal to the number of wing panels used in the UVLM. Any modes associated with frequencies at least ten times higher than the flapping frequency are neglected, as the modal contribution is expected to be negligible. The second natural frequency was 360 Hz for a homogeneous wing with elastic modulus of 10 GPa. This is roughly 14 times greater than the flapping frequency of 25 Hz. All other stiffness distributions tested have a greater second natural frequency, so only the first mode was retained for simulations.

**Table 1:** Discretization Parameters

| Symbol | Description | Value |
| --- | --- | --- |
| $N_e$ | Number of elements | 40 |
| $N_p$ | Number of panels | 40 |
| $N_w$ | Number of wake vortices | 100 |
| $T/dt$ | Time-steps per cycle | 200 |

We use an explicit solver that remains stable for relatively coarse time steps. Using a homogeneous wing, the model is stable at 22 time steps per wingbeat. When the taper ratio is reduced to 0.2, the solver required 100 timesteps per wingbeat. At the beginning of each time step, the deformed shape of the wing is used to calculate the aerodynamic influence coefficients. The inlet velocity is then calculated using the wake-induced velocity, wing deformation rate, and prescribed kinematics. Enforcing the non-penetration condition, the strengths of the wing-bound vortices are found via Eq. 19. The advection of the wake is calculated next, followed by the aerodynamic loads. These forces are subsequently used in Eq. 15 to determine the generalized load. Equation 12 is integrated using the MATLAB differential equation solver ‘ode45’. Lastly, the power consumption and reaction force are computed before moving to the next time step.

Wing morphology and flapping kinematics are based approximately on the *M. Sexta* [25, 8, 26] and are summarized in Tab. 2. Flapping kinematics assume hovering flight. Pitch and plunge are idealized as purely harmonic with a frequency of 25 Hz and a phase difference of −90°. The plunge amplitude was approximated by the distance traveled by the semispan (x = span/2) of a three-dimensional wing experiencing a flap amplitude of 60°. The Young’s modulus and density were estimated from measurements on insect cuticle [27, 28].

**Table 2:** Wing Parameters

| Symbol | Description | Value | Units |
| --- | --- | --- | --- |
| $E$ | Young’s Modulus | 10 | GPa |
| $c$ | Chord Length | 2 | cm |
| $b$ | Span | 5 | cm |
| $m$ | Mass | 45 | mg |
| $\zeta_1$ | First Modal Damping Ratio | 0.05 | - |
| $Z_0$ | Plunge Amplitude | 2.62 | cm |
| $\theta_0$ | Pitch Amplitude | 45 | $^\circ$ |
| $\beta_z$ | Pitch-Plunge Phase Difference | -90 | $^\circ$ |
| $\omega$ | Flapping Frequency | 25 | Hz |

## 4. Results

In this section, we show how taper ratio influences the dynamics of flapping wings. First, we determine how taper affects the wing’s first natural frequency and mode shape. Next, we calculate wingtip deflection, lift, power consumption and total forces acting at the wing’s leading edge for a wing flapping with hovering kinematics. We then conduct a parameter study to examine how deviations from hovering kinematics affect wing performance.

### 4.1. Natural Frequencies & Mode Shapes

The wing’s natural frequency affects deflection magnitude and consequently, the aerodynamic forces that arise from flapping. Certain flapping-to-natural frequency ratios have been shown to improve the aerodynamic and energetic performance of flapping wings [2, 5, 6]. Here, we explore how variable flexural rigidity influences the wing’s natural frequency to provide context to the dynamic studies described in the following sections.

Over the range of taper ratios considered, the wing’s natural frequency varies from about 57 to 115 Hz (Fig. 4). Assuming a flap frequency of 25 Hz, this coincides to a flapping-to-natural frequency ratio range from 0.21 to 0.44. Natural frequencies corresponding to the first torsional mode in *M. sexta* forewings have been reported between 75-95 Hz [26, 29], which are represented by taper ratios from about 0.4 to 0.5. Note that the first mode in the *M. sexta* forewing corresponds to a bending mode, however since our 2D model does not account for spanwise deformation, it is more appropriate to compare our computed natural frequencies to torsional modes where deformation varies predominately over the wing chord. Though the bending mode in *M. sexta* wings has a lower natural frequency (≈ 60 - 70 Hz,[26, 29]) than the twisting mode, this does not necessarily imply the bending mode is more important. The insect flaps at about 25 Hz, and previous studies have shown flapping excites frequencies at odd multiples of the wingbeat frequency. Consequently, twisting may be as significant as bending to wing deformation.

**Figure 4:**
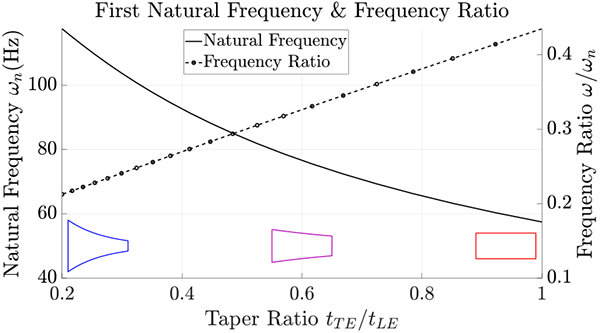
First natural frequency (solid line) and frequency ratio (dashed line) as a function of wing taper. Approximate airfoils indicating the taper are shown at the bottom of the figure, with taper ratios of 0.2, 0.6, and 1 from left to right. Flapping frequency is assumed to be 25 Hz.

**Figure 5:**
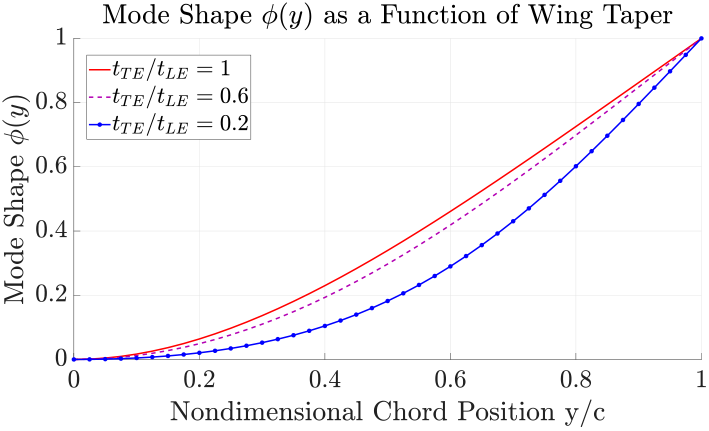
The bending mode used to calculate wing deflection. Mode shapes presented in this plot are normalized by the maximum value at the trailing edge.

As the taper ratio decreases from 1 to 0.2, the wing’s local area moment of inertia and mass increase near the leading edge and decrease towards the trailing edge. As a result, the wing’s natural frequency is inversely proportional to taper ratio (Fig. 4). The most tapered wing has a natural frequency nearly double that of the homogeneous wing. Mode shapes associated with tapered wings tend to bend more around the center of the chord, while bending is localized towards the leading edge in homogeneous wings. This is consistent with torsional mode shapes measured in *M. sexta* forewings [26].

### 4.2. Flapping With Hovering Kinematics

We now examine the wing’s dynamic response assuming the flapping kinematics described in Tab 2. Wing deflection, lift, power and leading edge force are shown for various taper ratios in Fig. 6. Quantities are reported as a function of wingbeat fraction *t/T*, where t is time and *T* is the wingbeat period. All quantities are taken once the wing has achieved steady-state. We use mean power as a proxy for energy consumption. This assumes negative power is elastically stored in the insect exoskeleton and that this stored potential energy can be recycled to offset positive power requirements. Then, we use the *Z* component of total leading edge force **F**_*T,XYZ*_ to approximate the forces the insect flight muscle must produce in order to flap the wing. **F**_*T,XYZ*_ is defined with respect to the inertial coordinate system and is inclusive of both aerodynamic and inertial forces. The *Z* component of **F**_*T,XYZ*_ (simply leading edge force hereafter) is a better proxy for muscle forces than total moment **M**_*T*_, since the literature suggests wing pitching arises passively from inertial and aerodynamic forces [30].

**Figure 6:**
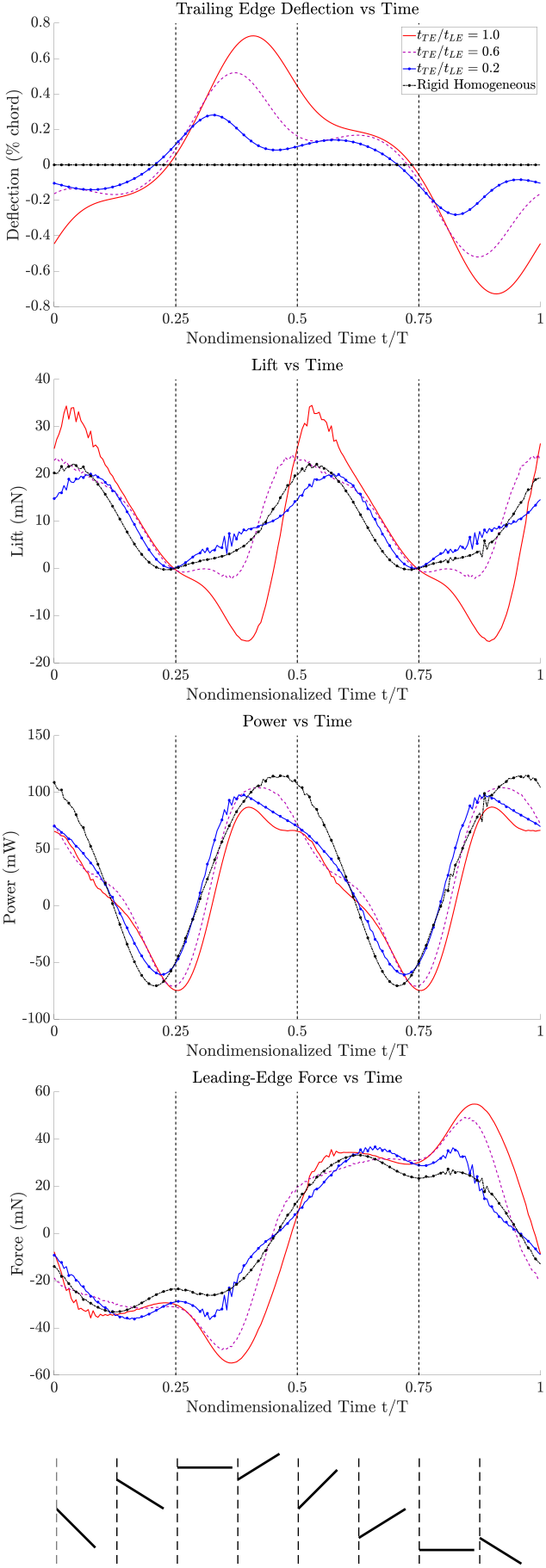
Comparison of lift, power, leading edge force, and wing tip deflection for several taper ratios using kinematics in Table 2. The bottom image indicates the pitch and plunge of the wing as a function of nondimensionalized time.

Calculated deflections appear to be consistent with reported values for *M. sexta* [25]. Previous studies reported the angle of rotation between the leading edge and the trailing edge, which is an aggregate quantity influenced both by the rigid body pitch angle and trailing edge deflection. Angles of rotation were typically around 45° for chords proximal to the insect body and 65° for chords distal to the insect body. Our model predicted angles of rotation of approximately 59° when for a taper ratio of 0.2 and 71° for a taper ratio of 1. While the homogeneous wing resulted in an angle of rotation greater than reported values, the tapered wings matched fairly well. Aerodynamic forces are also consistent with those produced by *M. sexta*. Assuming the insect has a body mass of 2.0 grams [31], a wing pair produces sufficient lift to hover when taper ratios are between about 0.27 and 0.65. This corresponds to a natural frequency range of about 70 to 109 Hz.

As wing taper increases, both wingtip deflection amplitude and primary response phase (taken with respect to plunge) decrease (Fig. 6). The lower deflection is due to the tapered wing being stiffer than the homogeneous wing, evidenced by the higher natural frequency (Fig. 4). Peak deflection moves away from the mid-stroke (*t/T* = 0, 0.5) and towards the stroke reversals (*t/T* = 0.25, 0.75) as wing flexibility increases. This indicates that deflection in highly tapered wings is more influenced by the plunging inertial force and less by the aerodynamics, as plunging acceleration *Z* is at a maximum while the aerodynamic force is minimized at stroke reversal.

Wing taper has a large effect on lift production. While the homogeneous wing generated the highest peak lift, its time-averaged lift was 20 to 30 percent lower than that of the other wings (Tab. 3). This can be partially attributed to the large troughs in lift occurring at *t/T* ≈ 0.375, 0.875, when the plunging motion of the wing reverses. These troughs are not seen with the stiffer wings. The lowest point in the trough roughly corresponds to the peak deflection experienced by the wing. Separating lift into steady and unsteady components, we see that the negative lift is largely driven by the steady component. When the wing undergoes large deflections, it reduces the effective angle of attack and creates an adverse camber that biases lift in the positive *Y* direction. As the taper becomes more pronounced and wing stiffness increases, the lift contribution from the steady component increases while that of the unsteady component is reduced.

**Table 3:** Peak deflection, mean lift, mean power and peak leading edge force for various taper ratios assuming normal flapping kinematics. Rigid wing is assumed to have uniform thickness.

| **Taper Ratio** | 1.0 | 0.6 | 0.2 | Rigid |
| --- | --- | --- | --- | --- |
| Peak deflection (mm) | 14.6 | 10.4 | 5.6 | 0 |
| Mean lift (mN) | 7.19 | 10.1 | 9.47 | 8.89 |
| Power (mW) | 15.5 | 26.4 | 28.8 | 33.6 |
| Peak leading edge force (mN) | 54.8 | 49.2 | 36.7 | 33.2 |

Mean power is sensitive to taper ratio as well. The primary effect of wing deformation is seen at the mid-stroke (*t/T* = 0, 0.5), where peak power is lower for the flexible wings compared to the rigid wing. At that instant, the flexible wings are bent backwards due to aerodynamic forces, and their angle of attack is lower compared to that of the rigid wing. As a result, mean power is lower for the flexible wing compared to the rigid wing (Tab. 3). While mean power reduces with taper ratio, peak leading edge forces increase. The most flexible wings experience the largest leading edge forces, where the peaks in forces coincide approximately with peak deflections. This indicates that the inertial and aerodynamic forces associated with elastic deformation increase the leading edge forces required to sustain the prescribed kinematics of *O*.

### 4.3. Deviations from Hovering Kinematics

In this section, we explore the sensitivity of the wing response to kinematic deviations from the hovering flapping kinematics shown in Tab. 2. We characterize the influence of taper ratio on lift, power and leading edge force for various sets of flapping kinematics.

#### 4.3.1. Pitch Amplitude

First, we investigate how pitch amplitude affects wing dynamics. Despite widespread usage as a kinematic variable, rigid-body pitch angle is difficult to characterize in flexible insect wings since chord-wise bending may affect how the angle is calculated from physical measurements. We tested several values of rigid-body pitch amplitude to examine how wing performance is affected by variations in this quantity (Fig. 7).

**Figure 7:**
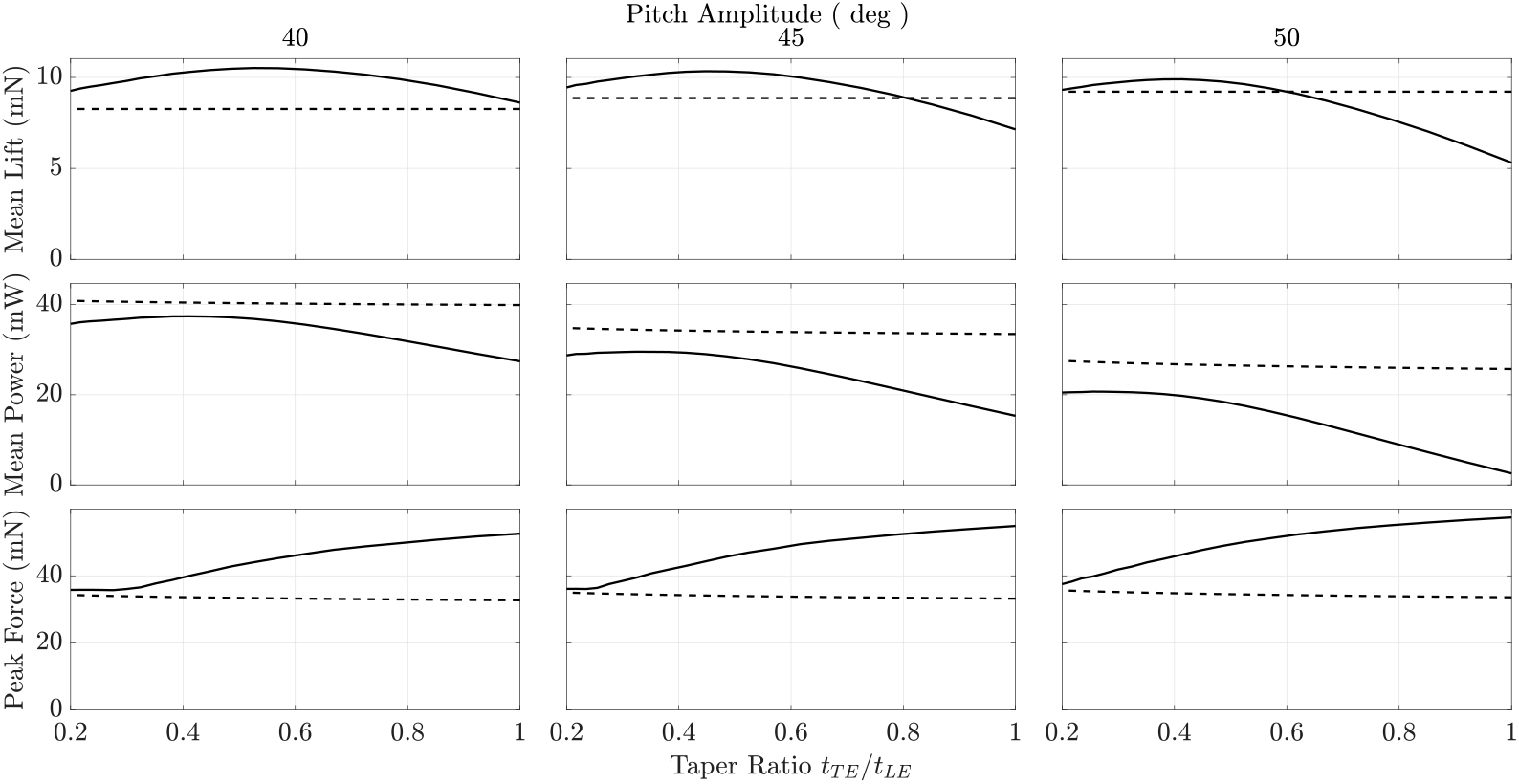
Study of pitch amplitude. The flexible and rigid wing values are represented by the solid and dashed lines, respectively.

**Figure 8:**
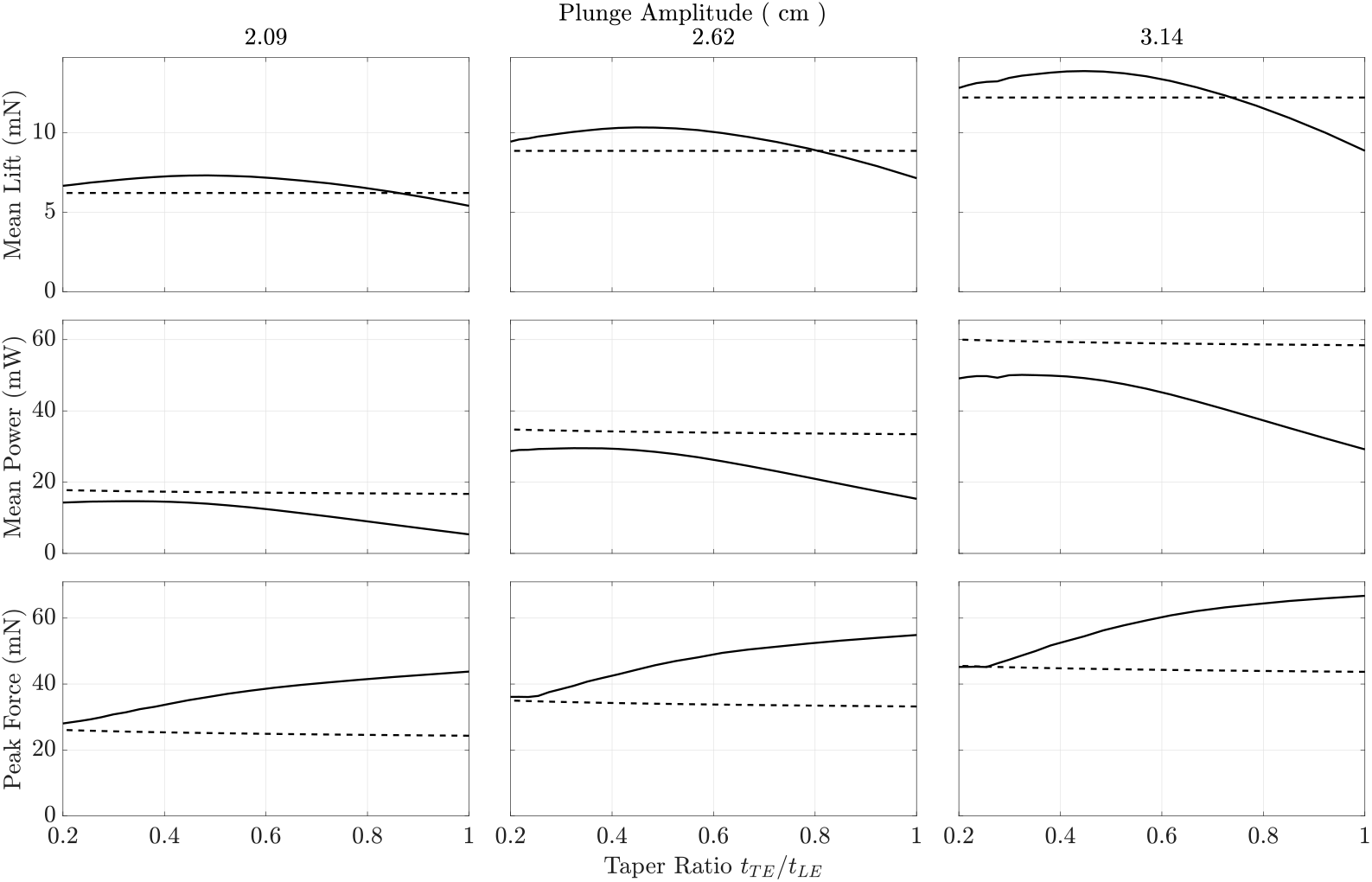
Study of plunge amplitude. The flexible and rigid wing values are represented by the solid and dashed lines, respectively.

Lift and power consumption are both strongly impacted by pitch amplitude. Pitch amplitude significantly affects mean lift in highly flexible wings, but only modestly affects lift in stiffer wings. When highly tapered, the wing produced nearly constant lift regardless of the pitching amplitude. For example, for a taper ratio of 0.2, the wing produces 9.26 mN for a pitch amplitude of 40° and to 9.45 mN for a pitch angle of 50°. By contrast, the highly flexible homogeneous wing is more sensitive to pitch amplitude. It generates less lift with increasing pitch angle, falling from 8.6 mN at 40° to 5.3 mN at 50°, a drop of nearly 40%. The wing experiences similar deformation under both pitch angles, but the effective angle of attack is decreased by the higher pitch angle in addition to the deflection-induced camber and the downwash caused by the wake vortices. Around stroke reversal in the 50° pitching case, the aerodynamic force is heavily weighted towards the leading edge, reducing both lift and power consumption. A byproduct of this is that the taper ratio associated with maximum lift decreases with increasing pitch amplitude, from 0.53 at 40° to 0.42 at 50°. These results also demonstrate that there is an optimal taper ratio which maximizes lift generation, and that this optimal taper ratio depends on pitch amplitude.

Power consumption of the flexible wing is lower than that of the rigid wing in all cases. The mean power versus taper ratio curve is concave, with the location of the maxima contingent on pitch amplitude. The low energy requirement of the flexible wing at high pitch amplitude is due to the wing bending out of the way and minimizing aerodynamic loads, though this adversely affects lift generation.

Peak leading edge force is relatively unaffected by the pitching angle. Forces in general are dominated by the plunging inertial and aerodynamic forces rather than the pitching inertial force. The pitching motion accounts for only 19% of the peak leading edge force in a homogeneous flexible wing and 14% in a wing with a taper ratio of 0.2. On the other hand, peak leading edge force is sensitive to taper ratio. The lowest leading edge forces occur in the most tapered wing, presumably because the wing’s center of mass is closer to the point of rotation. This reduces the inertial forces associated with pitching. The largest leading edge forces occur in the untapered wing, since the center of mass is further from the point of rotation and the wing experiences larger deflections.

#### 4.3.2. Plunge Amplitude

Some insects have been shown to increase their flapping amplitude to increase lift generation [32]. Within the pitch-plunge model, this is akin to increasing the plunge amplitude. We tested three different values of plunge amplitude to identify how variation of this parameter affected lift, mean power and peak leading edge forces (Fig. 7).

Plunge amplitude impacts each quantity strongly. Raising the plunge amplitude increases the distance and velocity of the wing during each stroke. As expected, this increases the lift, power, and leading edge force. Mean lift in flexible wings does not increase proportionally to the plunge velocity squared as might be expected from conventional steady aerodynamic theory. A possible reason for this is that the increased velocity generates higher aerodynamic forces on the wing, which in turn leads to higher deflections and reduced angle of attack, meaning that increasing the plunge amplitude and velocity results in diminishing returns. Mean lift in stiffer wings more closely scales with plunge velocity squared.

For a stationary airfoil subject to a uniform free-stream velocity, aerodynamic power scales with the free-stream velocity cubed [33]. We observe that the mean power scales nearly cubically with plunge velocity in rigid wings. In the highly tapered wing, the mean power requirement is nearly identical to that of a rigid wing at low plunge amplitudes. However, at higher plunge amplitudes, the disparity between power requirements is larger. The increased deflection arising from plunge amplitude likely decreases the effective drag on the wing, thereby reducing the power required by the flexible wing. This also explains why power increases with taper ratio for all plunge amplitudes considered, though mean lift decreases above a certain taper ratio as well. The optimal taper ratio thus falls somewhere between providing sufficient lift to fly while requiring low energetic expenditures. Unlike pitch amplitude, the plunge amplitude does not affect the taper ratio associated with peak lift.

Increasing the plunge amplitude increases the peak leading edge force for all wings considered, since the increased plunging amplitudes lead to larger linear accelerations and aerodynamic forces. As plunge amplitude increases, the contribution of the plunging force actually decreases slightly for all tested taper ratios. This is because the aerodynamic forces acting on the wing are more sensitive to increases in plunging amplitude than the plunging inertial forces are.

#### 4.3.3. Pitch-Plunge Phase

Finally, we explore how the relative phase between pitching and plunging affects flapping wing aeromechanics. Hawkmoths have been shown to adjust the phase of their muscle activation in response to visual cues that simulate upward or downward motion [34]. Based on this evidence, it is plausible that modulation of the phase between kinematic parameters is responsible for some maneuvers. Mean lift, power and peak leading edge forces are shown as a function of taper ratio for various pitch-plunge phase differences in Fig. 9.

**Figure 9:**
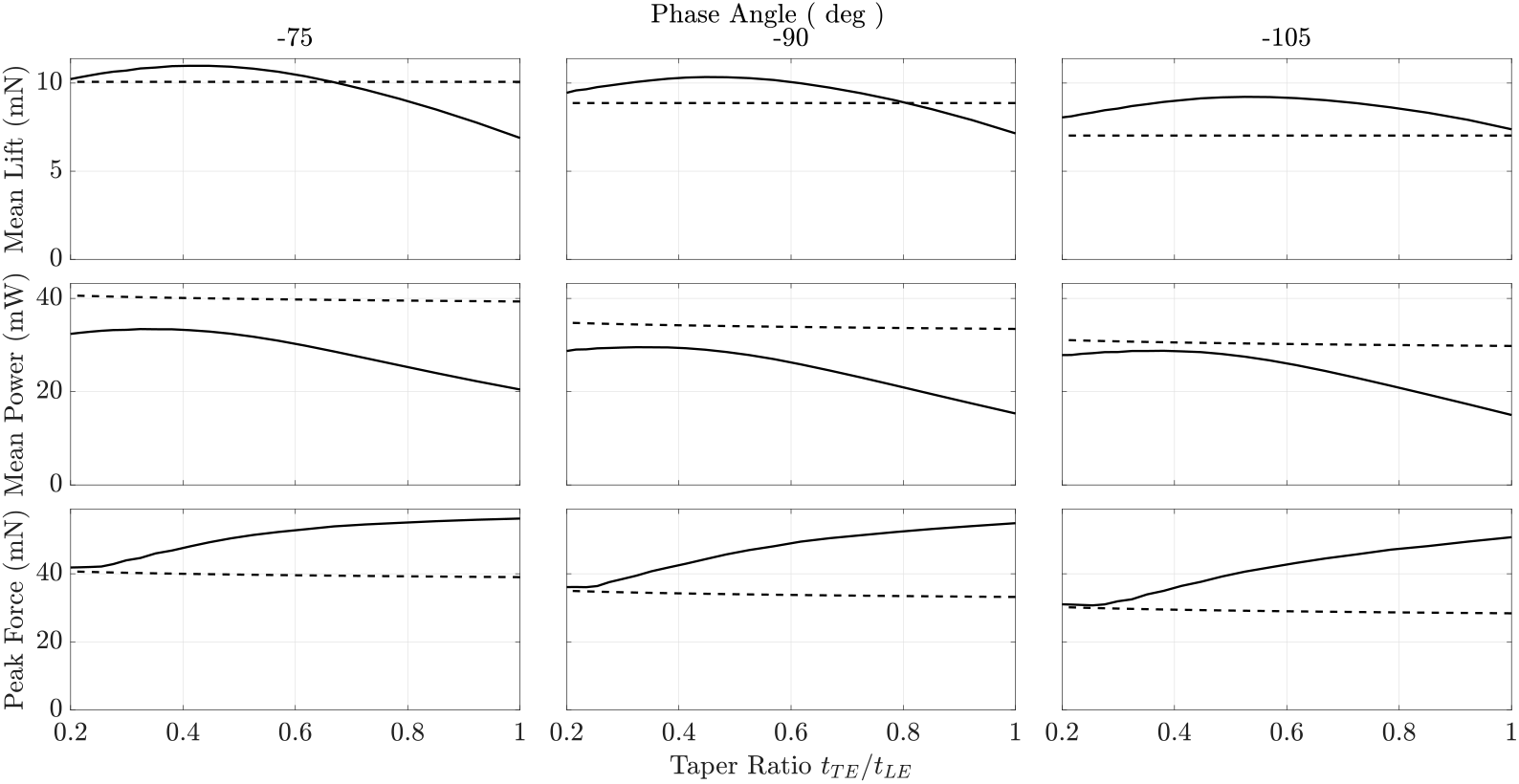
Study of pitch-plunge phase offset. The flexible and rigid wing values are represented by the solid and dashed lines, respectively.

Mean lift is fairly influenced by deviations in pitch-plunge phase. Increasing the phase to −75° increases the highest achievable lift in both flexible and rigid wings, whereas decreasing the phase to −105° decreases highest achievable lift. Flexible wings across all taper ratios produce higher lift than rigid wings for a pitch-plunge phase of −105°, but for a pitch-plunge phase of −75°, flexible wings only produce mean lift than rigid wings for low taper ratios. Similar to trends observed in pitch amplitude, the taper ratio associated with greatest mean lift varies modestly with respect to pitch-plunge phase.

Mean power in general decreases when pitch-plunge phase is lowered. The largest reduction in mean power occurs in the rigid wing. The flexible wing have similar power requirements for −105° and −90° pitch-plunge phases, while power requirements for −75° are slightly higher, presumably due to an increase in lift and drag. The opposite trends in leading edge force are true, where leading edge force is reduced as the phase angle is decreased. As with all other parameter studies considered, the peak leading edge force increases with taper ratio as the wing becomes more flexible.

## 5. Discussion

### 5.1. Optimal Lift Production

Functionally graded flexural rigidity may improve flight performance in terms of aerodynamic force generation across a broad range of flapping wing kinematics. Figures 7-9 show that the mean lift produced by flexible wings is generally greater than that produced by rigid wings, though rigid wings may outperform flexible wings when deformation is extreme. Interestingly, there is an optimal taper ratio that maximizes the mean lift. For the normal flapping kinematics shown in Tab. 2, this taper ratio is about 0.5, which corresponds to a natural frequency of 83 Hz and a flapping-to-natural frequency ratio of about 0.3. Prior studies show similar aerodynamic benefits when operating close to this ratio. Vanella et al. showed that flapping at 1/3 the wing’s natural frequency evokes a super-harmonic resonant response that improves the lift-to-drag ratio compared to a rigid wing with identical kinematics [2]. Dai et al. found that flexible wings with flapping-to-natural frequency ratios equal to or less than 0.3 produced more lift than rigid wings [35]. Though our results generally agree with these findings, we saw that the exact lift-maximizing flapping-to-natural frequency ratio depends on flapping kinematics. For a pitching amplitude of 50°, the optimal flapping-to-natural frequency ratio is about 0.27; this increases to about 0.35 when the pitching amplitude is reduced to 40°. The optimal flapping-to-natural ratio changes with pitch-plunge phase as well, but is stationary with respect to plunge amplitude. This collectively shows that the wing’s natural frequency is strongly associated with lift generation.

### 5.2. Trade-offs Between Power & Force

In all cases tested, mean power and peak leading edge force were inversely proportional. With increasing taper ratio, mean power decreased while peak leading edge force increased. As discussed previously, the reduction of power at high taper ratios likely stems from a reduction of drag, though lift is lower at higher taper ratios as well. Large leading edge forces at high taper ratios result from the increased deflections and accompanying inertial forces.

The inverse relationship of mean power and peak leading edge force poses trade-offs for flight. Low mean power consumption implies the insect requires less energy storage, and low peak leading edge forces implies that the insect requires less muscle mass. Based on insect anatomy, we hypothesize that the flight mechanism is oriented more towards energetic efficiency. The thorax constitutes about 25-30% of the total body mass in *M. sexta* [31] and the flight muscles occupy much of the thorax volume. The flight muscles can generate staggering forces, with one flight muscle group producing forces as high as 1 N [36]. These forces are 50 times greater than the total weight of the insect assuming a body mass of 2 grams. It is possible that the increased muscle force requirements resulting from wing flexibility are an acceptable trade-off for high aerodynamic force generation and relatively low power consumption.

### 5.3. Computational Economy

The reduced-order FSI model developed through this work can be used to estimate how morphological or material properties beyond flexural rigidity impact flapping wing performance. Though the pitch-plunge idealization is used frequently to study of flapping wings, most implementations still rely on high-order computational solvers. UVLM has been incorporated into the pitch-plunge model to reduce the computational costs of the fluid solver on several occasions, but the AMM representation is less frequently used to lower the costs of the structural solver. Together, our model based on coupled UVLM/AMM can be solved in less than one second per wingbeat for the described discretization parameters, representing a large computational savings compared to higher-order solvers. When running on a laptop with a Ryzen 5800H 3.2 GHz CPU and 16 GB of DDR4 RAM, our model solves in roughly 10 seconds when considering 20 wing beat cycles. Further, to our knowledge, the flapping wing moments, leading edge forces and power for a flexible wing experiencing pitch-plunge motion have not previously been derived in modal coordinates. If wing deformation is known, these expressions can be used to compute quantities more efficiently than if using the canonical definitions, since spatial and temporal dependence can be decoupled and spatial integrals pre-computed.

While computationally efficient, our FSI model has limitations as well. First, the simplified 2D model neglects some dynamics associated with 3D flapping, such as spanwise bending, spanwise flow and wingtip losses. Previous studies have shown that spanwise flow may positively or negatively influence lift depending on the relative phase between bending and flapping [18]. Spanwise bending may be incorporated in future work and would best be accounted for by modifying plunge to be the sum of rigid body flapping and elastic deformation, where the elastic deformation is a degree-of-freedom to be solved for. Spanwise flow and wingtip losses are more difficult to accommodate in a 2D setting, and consequently 2D models are most appropriate to generate approximate low-fidelity solutions quickly. The validity of the 2D solution, and how spanwise flow and wingtip loss may affect a specific wing configuration, should subsequently be assessed with higher-fidelity 3D modeling. Nonetheless, in many cases results from 2D have agreed with results generated by computational or experimental 3D studies with reasonable accuracy [37–39]. This suggests that at least some of the insights garnered from a 2D study can inform a 3D simulation. Next, the UVLM fluid model neglects viscous effects. While the inviscid assumption is appropriate for a broad range of flying insects including *M. sexta*, fluid viscosity may play a non-trivial role in the flight of very small insects [40]. Finally, the structural model is based on linear assumptions and cannot account for structural non-linearity. As a result, some dynamic phenomena associated with large deformation may be lost. Further, numerous features observed in real wing structures, such as venation patterns, curvature, corrugation and graded material properties are neglected. Each of these may contribute to the wing’s overall flexural rigidity, but in the authors’ opinion, must first be studied in isolation before a high-fidelity structural model of the wing is attainable.

## 6. Conclusion

Flapping insect wings are complex structures that deform from aerodynamic and inertial forces. In addition to the flapping motions of the wing, heterogeneous geometric and material properties influence how the wing deforms. Previous studies have shown that flexural rigidity in some insect wings varies as much as two orders of magnitude from the leading edge to the trailing edge [8]. However, the computational cost of many flapping wing FSI models have precluded us from understanding how these gradients in flexural rigidity regulate flapping wing performance. In this work, we developed a simplified 2D FSI model of a flapping wing as a pitching-plunging airfoil using AMM and UVLM. We derived expressions for the power, leading edge forces and moments required to flap the wing using model coordinates. We used these models to understand how gradients in flexural rigidity affect flight energetics and the forces needed to sustain flight.

We found that wings with functionally graded flexural rigidity increased aerodynamic force generation and decreased power requirements relative to rigid wings or wings with uniform flexural rigidity. For normal hovering flapping kinematics, a wing with an optimal taper ratio of 0.5 produced about 20% more mean lift than a rigid wing and required about 15% less mean power. This taper ratio corresponded to a wing with a natural frequency of about 85 Hz, which is similar to the torsional natural frequency measured in *M. sexta* wings [26]. Variation of rigid body pitch amplitude and relative pitch-plunge phase affected the taper ratio that maximized lift production. Otherwise, trends in mean lift, mean power and peak leading edge force were not dramatically affected by deviations in flapping kinematics. This work suggests that functionally graded flexural rigidity, driven by spatial variations of wing thickness, enhances the performance of flapping wing insects.

## Acknowledgment

This research was supported the National Science Foundation under awards Nos. CBET-1855383 and CMMI-1942810 to MJ. Any opinions, findings, and conclusions or recommendations expressed in this material are those of the author(s) and do not necessarily reflect the views of the National Science Foundation.

## References

[1] S. A. Combes, T. L. Daniel, Into thin air: contributions of aerodynamic and inertial-elastic forces to wing bending in the hawkmoth manduca sexta, Journal of Experimental Biology 206 (2003) 2999–3006.

[2] M. Vanella, T. Fitzgerald, S. Preidikman, E. Balaras, B. Balachandran, Influence of flexibility on the aerodynamic performance of a hovering wing, Journal of Experimental Biology 212 (2009) 95–105.

[3] A. Mountcastle, T. Daniel, Vortexlet models of flapping flexible wings show tuning for force production and control, Bioinspiration & biomimetics 5 (2010) 045005.

[4] A. M. Mountcastle, S. A. Combes, Wing flexibility enhances load-lifting capacity in bumblebees, Proceedings of the Royal Society B: Biological Sciences 280 (2013) 20130531.

[5] M. Jankauski, Z. Guo, I. Shen, The effect of structural deformation on flapping wing energetics, Journal of Sound and Vibration 429 (2018) 176–192.

[6] H. E. Reid, R. K. Schwab, M. Maxcer, R. K. Peterson, E. L. Johnson, M. Jankauski, Wing flexibility reduces the energetic requirements of insect flight, Bioinspiration & biomimetics 14 (2019) 056007.

[7] B. Yin, H. Luo, Effect of wing inertia on hovering performance of flexible flapping wings, Physics of Fluids 22 (2010) 111902.

[8] S. Combes, T. Daniel, Flexural stiffness in insect wings ii. spatial distribution and dynamic wing bending, Journal of Experimental Biology 206 (2003) 2989–2997.

[9] F.-B. Tian, H. Dai, H. Luo, J. F. Doyle, B. Rousseau, Fluid–structure interaction involving large deformations: 3d simulations and applications to biological systems, Journal of computational physics 258 (2014) 451–469.

[10] D. Ishihara, T. Horie, M. Denda, A two-dimensional computational study on the fluid–structure interaction cause of wing pitch changes in dipteran flapping flight, Journal of Experimental Biology 212 (2009) 1–10.

[11] M. Hamamoto, Y. Ohta, K. Hara, T. Hisada, Application of fluid–structure interaction analysis to flapping flight of insects with deformable wings, Advanced Robotics 21 (2007) 1–21.

[12] T. Fitzgerald, M. Valdez, M. Vanella, E. Balaras, B. Balachandran, Flexible flapping systems: Computational investigations into fluid-structure interactions, The Aeronautical Journal 115 (2011) 593–604.

[13] J. Tang, S. Chimakurthi, R. Palacios, C. Cesnik, W. Shyy, Computational fluid-structure interaction of a deformable flapping wing for micro air vehicle applications, in: 46th AIAA Aerospace Sciences Meeting and Exhibit, 2008, p. 615.

[14] M. Jankauski, I. Shen, Dynamic modeling of an insect wing subject to three-dimensional rotation, International Journal of Micro Air Vehicles 6 (2014) 231–251.

[15] J. D. Anderson, J. Wendt, Computational fluid dynamics, volume 206, Springer, 1995.

[16] M. F. Platzer, K. D. Jones, J. Young, J. C. Lai, Flapping wing aerodynamics: progress and challenges, AIAA journal 46 (2008) 2136–2149.

[17] F.-B. Tian, H. Luo, J. Song, X.-Y. Lu, Force production and asymmetric deformation of a flexible flapping wing in forward flight, Journal of Fluids and Structures 36 (2013) 149–161.

[18] B. A. Roccia, S. Preidikman, M. L. Verstraete, D. T. Mook, Influence of spanwise twisting and bending on lift generation in mav-like flapping wings, Journal of Aerospace Engineering 30 (2017) 04016079.

[19] J. M. Birch, M. H. Dickinson, Spanwise flow and the attachment of the leading-edge vortex on insect wings, Nature 412 (2001) 729–733.

[20] J. Kweon, H. Choi, Sectional lift coefficient of a flapping wing in hovering motion, Physics of Fluids 22 (2010) 071703.

[21] J. Hoffmann, S. Donoughe, K. Li, M. K. Salcedo, C. H. Rycroft, A simple developmental model recapitulates complex insect wing venation patterns, Proceedings of the National Academy of Sciences 115 (2018) 9905–9910.

[22] J. Reade, M. A. Jankauski, Deformable blade element and unsteady vortex lattice fluid-structure interaction modeling of a 2d flapping wing, in: International Design Engineering Technical Conferences and Computers and Information in Engineering Conference, volume 83969, American Society of Mechanical Engineers, 2020, p. V007T07A032.

[23] J. Katz, A. Plotkin, Low-speed aerodynamics, volume 13, Cambridge university press, 2001.

[24] L. Tang, M. P. Pai, et al., On the instability and the post-critical behaviour of two-dimensional cantilevered flexible plates in axial flow, Journal of Sound and Vibration 305 (2007) 97–115.

[25] A. P. Willmott, C. P. Ellington, The mechanics of flight in the hawkmoth manduca sexta. i. kinematics of hovering and forward flight., The Journal of experimental biology 200 (1997) 2705–2722.

[26] H. Reid, H. Zhou, M. Maxcer, R. K. Peterson, J. Deng, M. Jankauski, Toward the design of dynamically similar artificial insect wings, International Journal of Micro Air Vehicles 13 (2021) 1756829321992138.

[27] C. Casey, C. Yager, M. Jankauski, C. M. Heveran, The flying insect thoracic cuticle is heterogenous in structure and in thickness-dependent modulus gradation, bioRxiv (2021).

[28] J. Vincent, U. Wegst, Design and mechanical properties of insect cuticle., Arthropod structure & development 33 3 (2004) 187–99.

[29] A. G. Norris, A. N. Palazotto, R. G. Cobb, Experimental structural dynamic characterization of the hawkmoth (manduca sexta) forewing, International Journal of Micro Air Vehicles 5 (2013) 39–54.

[30] A. J. Bergou, S. Xu, Z. J. Wang, Passive wing pitch reversal in insect flight, Journal of Fluid Mechanics 591 (2007) 321–337.

[31] T. L. Hedrick, T. Daniel, Flight control in the hawkmoth manduca sexta: the inverse problem of hovering, Journal of Experimental Biology 209 (2006) 3114–3130.

[32] D. L. Altshuler, W. B. Dickson, J. T. Vance, S. P. Roberts, M. H. Dickinson, Short-amplitude high-frequency wing strokes determine the aerodynamics of honeybee flight, Proceedings of the National Academy of Sciences 102 (2005) 18213–18218.

[33] S. Schmitz, Aerodynamics of wind turbines: a physical basis for analysis and design, John wiley & sons, 2020.

[34] N. Ando, R. Kanzaki, Flexibility and control of thorax deformation during hawkmoth flight, Biology letters 12 (2016) 20150733.

[35] H. Dai, H. Luo, J. F. Doyle, Dynamic pitching of an elastic rectangular wing in hovering motion, Journal of Fluid Mechanics 693 (2012) 473–499.

[36] M. S. Tu, T. L. Daniel, Submaximal power output from the dorsolon-gitudinal flight muscles of the hawkmoth manduca sexta, Journal of Experimental Biology 207 (2004) 4651–4662.

[37] Z. J. Wang, J. M. Birch, M. H. Dickinson, Unsteady forces and flows in low reynolds number hovering flight: two-dimensional computations vs robotic wing experiments, Journal of Experimental Biology 207 (2004) 449–460.

[38] J. P. Whitney, R. J. Wood, Aeromechanics of passive rotation in flapping flight, Journal of Fluid Mechanics 660 (2010) 197–220.

[39] L. Liu, M. Sun, The added mass forces in insect flapping wings, Journal of theoretical biology 437 (2018) 45–50.

[40] A. Santhanakrishnan, A. K. Robinson, S. Jones, A. A. Low, S. Gadi, T. L. Hedrick, L. A. Miller, Clap and fling mechanism with interacting porous wings in tiny insect flight, Journal of Experimental Biology 217 (2014) 3898–3909.

